# *Tudor domain containing protein 5-like* (*Tdrd5l*) identifies a novel germline body and regulates maternal RNAs during oogenesis

**DOI:** 10.1101/2022.08.02.502436

**Authors:** Caitlin Pozmanter, Leif Benner, Sydney E Kelly, Harrison Curnutte, Mark Van Doren

## Abstract

Tudor domain-containing proteins are conserved across the animal kingdom for their function in germline development and fertility. Previously, we demonstrated that *Tudor domain-containing protein 5-like (Tdrd5l)* plays an important role in the germline where it promotes male identity. However, Tdrd5l is also expressed in both the ovary and testis during later stages of germline development, suggesting that it plays a role in germline differentiation in both sexes. We found that Tdrd5l localizes to a potentially novel germline body and plays a role in post-transcriptional gene regulation. RNA sequencing of *Tdrd5l*-mutant ovaries compared to wild-type showed that differentially expressed genes were enriched for maternally deposited RNAs. Additionally, embryos laid by *Tdrd5l*-mutant females exhibited reduced viability and displayed dorsal appendage defects suggesting a failure of proper dorsal-ventral (D/V) patterning. As D/V patterning is dependent on *gurken (grk)*, we examined Grk expression during oogenesis. We observed premature accumulation of Grk protein in nurse cells indicating that translation is no longer properly repressed during mRNA transport to the oocyte. We also observed increased nurse cell accumulation of the cytoplasmic polyadenylation element binding protein Oo18 RNA-Binding Protein (Orb or CPEB), a translational activator of Grk. Decreasing *orb* function was able to partially rescue the *Tdrd5l*-mutant phenotype, and so defects in Orb are likely a primary cause of the defects in *Tdrd5l* mutants. Our data indicate that *Tdrd5l* is important for translational repression of maternal mRNAs such as *orb*, and possibly others, following their synthesis in the nurse cells and during their transport to the oocyte.

## Introduction

Tudor domain-containing proteins are conserved across the animal kingdom and one important function of these proteins is germline specification and maintenance (1). Mutations in the genes for Tudor proteins have been shown to cause infertility in flies and mammals alike (2,3). While the molecular function of a tudor domain is to promote protein-protein interaction by binding to dimethylated arginine and lysine residues (4,5), many tudor domain-containing proteins also have additional domains such as RNA recognition motifs (6). Together this allows for tudor domains to scaffold larger protein and ribonucleoprotein complexes (RNPs) such as in RNA granules.

RNA granules allow for multiple modes of RNA regulation in locally concentrated and protected membrane-less organelles (7). One germline-specific granule that is enriched for tudor domain-containing proteins is the nuage (8–10), which produces piRNAs that silence transposons (11). In addition to the nuage, germ cells have a number of other types of granules that can also be found in other tissues, the most common of which are the processing bodies (P-bodies) (12). These granules can take on different functions all centered around post-transcriptional regulation of mRNAs such as inducing mRNA degradation via removal of the 5’ cap and translational repression via regulation of the poly(A) tail. mRNAs are often targeted to P-bodies through binding of miRNAs which leads to the formation of a protein complex around the RNA (13,14). RNA granules are particularly important in the germline where RNAs are often transcribed at a different time or place from their translation (15). mRNAs can be sent to these granules for repression until they are needed for translation or to form RNPs that are translationally repressed and targeted for transport to another location (16).

One special class of germline RNAs are the maternal RNAs generated during oogenesis and deposited into the oocyte for control of early embryonic development, in particular before activation of the zygotic genome (17). During *Drosophila* oogenesis, maternal RNAs are transcribed in nurse cells and then transported into the oocyte where they are later translated. While in the nurse cells, maternally deposited RNAs stay translationally silenced through mechanisms that are not totally understood (18). Some of these RNAs become localized to specific regions of the oocyte and act to establish the body axes of the embryo. One such RNA, *gurken (grk)*, is translated first at the posterior of the oocyte to specify posterior fate before localizing to the dorsal-anterior corner of the oocyte to specify dorsal fate (19–21). Other important RNAs include *oskar (osk)*, which localizes to the posterior to help specify posterior identity and the germline (22) (23), and *bicoid (bcd)*, which specifies the anterior (24).

Previous work in our lab identified a novel tudor domain-containing protein, Tdrd5l (Tudor domain-containing protein 5-like), during a screen for genes involved in germline sex determination (25). Tdrd5l is homologous to mouse Tdrd5 as well as to another Drosophila protein, Tejas, both of which are required for fertility (10,26). All of these proteins have a highly related tudor domain, while Tejas and mouse Tdrd5 also have LOTUS domains that interact with the RNA helicase Vasa (27). *Drosophila* Tejas and mouse Tdrd5 both localize with Vasa to the nuage where they act in piRNA synthesis and retrotransposon repression (10,26).

While Tdrd5l is male-specific in the early germline, it is expressed in differentiating germ cells in both males and females, suggesting that it plays a role in post-transcriptional gene regulation of both spermatogenesis and oogenesis. In this study we demonstrate that Tdrd5l localizes to a novel germline body that associates with, but is distinct from, the nuage. Further, we demonstrate that, in females, *Tdrd5l* is important for post-transcriptional regulation of maternal RNAs. Embryos from *Tdrd5l*-mutant mothers exhibit defects in dorsal-ventral patterning and both Grk and Orb (Oo-18 RNA-Binding Protein) prematurely accumulate in nurse cells instead of being primarily expressed in the oocyte. Thus, one important role for*Tdrd5l* is in regulation of maternal RNAs as they are being transported from the nurse cells to the oocyte.

## Results

### Tdrd5l localizes to cytoplasmic bodies in the *Drosophila* germline

Previously, we characterized Tdrd5l expression using an N-terminally tagged exogenous BAC construct (25). Subsequently, we have used CRISPR/CAS9 to generate both N-and C-terminally tagged versions of the endogenous *Tdrd5l* locus. Interestingly, both N-and C-terminally tagged alleles behaved as loss of function alleles, indicating that these regions are important for *Tdrd5l* function. In particular, the N-terminally tagged allele exhibited localization to punctae in the germline when wild-type (wt) Tdrd5l protein was present, but not when the N-terminally tagged allele was the only source of Tdrd5l protein (Fig S1). This indicates that the N-terminus is important for localization of the Tdrd5l protein, but that there is an independent mechanism for Tdrd5l localization to punctae dependent on the wt protein. To visualize localization of functional Tdrd5l protein, we generated internal hemagglutinin (HA) and FLAG epitope-tagged alleles (Fig S1A), which behaved as wt alleles of *Tdrd5l*. We also generated a peptide guinea pig polyclonal antibody using amino acids 65-81 of Tdrd5l. This antiserum yielded a similar staining pattern to the internally tagged *Tdrd5l* alleles and immunostaining was eliminated in *Tdrd5l* mutants (Figure S2).

Using the internally FLAG-tagged allele, we observed anti-FLAG immunoreactivity in a mixed population of small and larger cytoplasmic bodies in the pre-meiotic male germline (Fig 1A), in agreement with our previous work using the exogenous HA-tagged BAC construct (25). In the germline stem cells (GSCs) and gonioblasts (GBs) (Fig 1A white arrow) there was a trend to have more smaller bodies and the occasional large body. In later spermatogonia and spermatocytes the granules are almost all large bodies 0.9-1 micron in diameter (Fig 1A yellow arrow). Interestingly, in the larger bodies Tdrd5l localized in a hollow, spherical pattern indicating that it formed the periphery of the body (Fig 1G).

**Figure 1:**
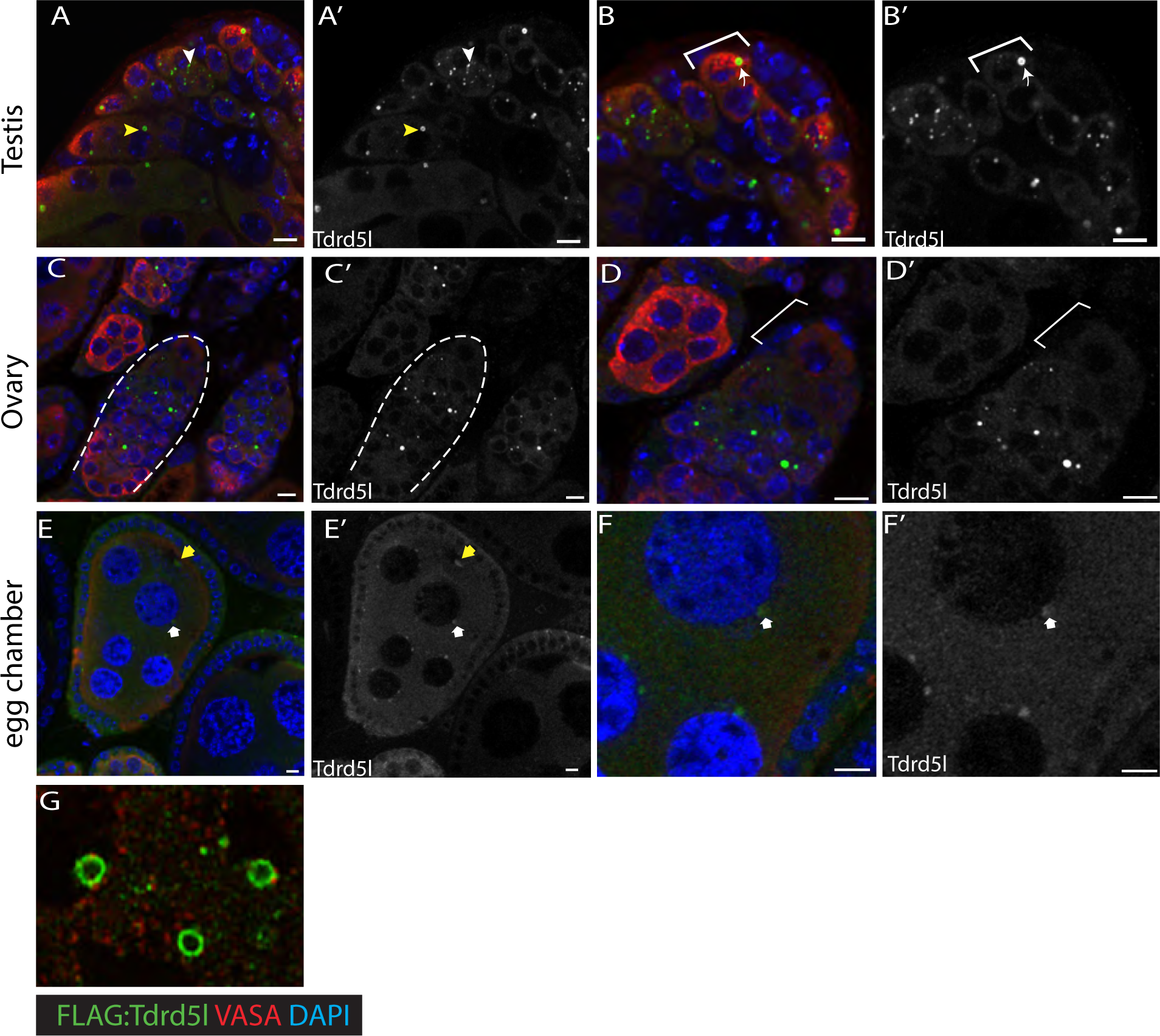
Tdrd5l localizes to cytoplasmic bodies in the *Drosophila* germline. (A-G) immunofluorescence of adult gonads using an endogenously FLAG-tagged allele of*Tdrd5l*. Panels B, D and F are higher magnification images of panels A, C and E, respectively. Antibodies used are as indicated in the figure. Scale bars = 5 microns. A,B) Adult testes. Note that Tdrd5l localizes to punctae or “bodies” in germ cells. Examples of small bodies are marked by white arrow heads and examples of large bodies are marked by yellow arrowheads. GSCs are indicated by brackets. C,D) Adult ovaries. A germarium is indicated by the dashed line in C. Tdrd5l localizes to bodies in the germline similar to those observed in testes but are not observed in female GSCs (brackets). E,F) Stage 7 egg chamber. Tdrd5l localizes to cytoplasmic bodies in both nurse cells (white arrow) and oocyte (yellow arrow). G) Immunofluorescence of an adult testis using an LSM 980 with an airyscan detector. Note the localization of FLAG-Tdrd5l to the periphery of the body.

In the female germline, Tdrd5l immunostaining was greatly reduced compared to the male germline when using identical confocal settings (Fig S3A-B), but still localized to cytoplasmic bodies (Fig 1C). Unlike in the male GSCs (Fig 1B brackets), Tdrd5l expression was absent from the germline stem cells (GSCs) in females (Fig 1D, brackets). Since Sxl represses Tdrd5l expression (25), this likely accounts for the lack of Tdrd5l expression in female GSCs where Sxl is expressed at high levels (28). At later stages in the female germline, we observed Tdrd5l localizing to bodies in the early differentiating germline of the germarium (Fig 1C), as well as in the nurse cells (Fig 1E and 1F, white arrow) and oocyte (Fig 1E, yellow arrow) until stages 6-7 of oogenesis.

### Tdrd5l localizes to an unknown body

Since many tudor domain-containing proteins, including the Tdrd5l homolog Tejas, are known to localize to RNA granules, we next analyzed the “Tdrd5l Body” to determine whether it corresponds to any previously identified RNA granules.

One prominent germline RNA granule is the perinuclear nuage, which acts as the site of piRNA production for transposon repression (29–31) and contains other tudor-domain containing proteins such as the Tdrd5l homolog Tejas in flies, and TDRD5 in mice (8,9,32). One hallmark of the nuage is the presence of Vasa protein (33). We observed that the Tdrd5l-containing bodies often associate with, or are imbedded in regions of Vasa immunoreactivity, but that Vasa staining is not detected within the Tdrd5l bodies themselves (Fig 2A, yellow arrow). Analysis of the distribution of Tdrd5l bodies revealed that there was an equal number that exhibited cytoplasmic vs. perinuclear localization, and bodies in both locations were associated with Vasa immunoreactivity (Fig S4A-B, N= 236 granules). Additionally, we co-stained for Tdrd5l and either Ago3 (Fig 2C, arrows) or Aub (Fig 2B, arrows), two other components of the nuage that are involved in piRNA biogenesis. In both cases we saw similar results as with Vasa, with the proteins enriched next to but not within the Tdrd5l body. These data indicate that either the Tdrd5l body is distinct from the nuage or forms a sub-compartment of nuage not marked by piRNA biogenesis factors. Consistent with this, we observed no changes in transposon expression in *Tdrd5l* mutant gonads of both sexes ((25) and Fig S5), thus suggesting a function for Tdrd5l distinct from its homolog Tejas.

**Figure 2:**
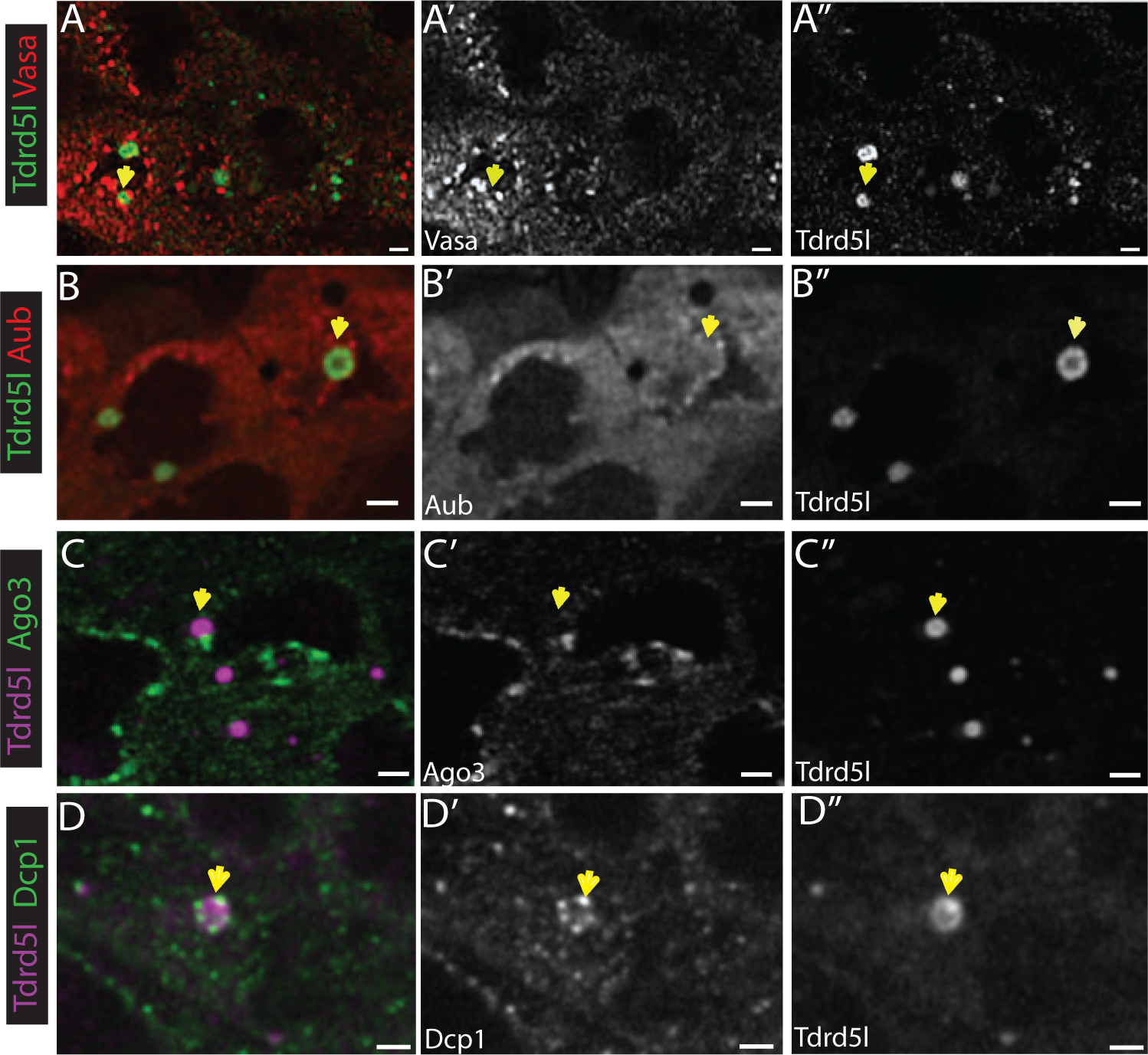
Tdrd5l localizes to a novel germline body. (A-D) Immunofluorescence of adult testes using endogenously FLAG-tagged *Tdrd5l* compared to markers for known RNA granules as indicated in figure. Examples of Tdrd5l bodies are marked with yellow arrows. A) Immunostaining of gonads for Vasa-positive nuage shows that Vasa staining is observed adjacent to, but not within, Tdrd5l bodies. Scale bar = 0.5 micron. (B,C) Immunostaining for Aub (B) or Ago3 (C) and Tdrd5l shows a similar relationship as Tdrd5l and Vasa, with Nuage factors being adjacent to, but not within, Tdrd5l bodies. D) Immunostaining of endogenously YFP-tagged Dcp1 and Tdrd5l bodies similarly shows punctae of Dcp1-YFP adjacent to, but not within, Tdrd5l bodies. Scale bars in B-D equal 1 micron.

Two additional RNA granules that are commonly found in many cell types are the P bodies, sites of post-transcriptional RNA regulation, and U bodies, which are involved in snRNP maturation (34). A common P body component is Decapping protein 1 (Dcp1) (35) and we visualized Dcp1 immunofluorescence in combination with Tdrd5l (Fig 2D). We did not observe co-localization between Dcp1-containing punctae and Tdrd5l, and the Dcp1-containing bodies were also of much smaller size than Tdrd5l bodies. In 63% of testes, we did observe Dcp1 labeled P-bodies associated with the periphery of Tdrd5l bodies (e.g. Fig 2D arrow, N= 17 testes). This suggests that there may be a relationship between P bodies and Tdrd5l bodies or exchange of materials between these structures, but these data indicate that the two types of bodies are distinct. We visualized U bodies using an endogenously tagged allele of Survival Motor Neuron (SMN) (34) and we failed to observe co-localization between SMN-containing bodies and Tdrd5l (Fig S4C,). Taken together, these data indicate that the Tdrd5l bodies are distinct structures from the Vasa positive nuage, P bodies and U bodies.

### Tdrd5l genetically interacts with post-transcriptional regulatory factors

Since cytoplasmic bodies are often important for post-transcriptional gene regulation, we determined whether *Tdrd5l* exhibited genetic interaction with known post-transcriptional regulation pathways. To ask whether Tdrd5l functions in regulating translation and RNA stability by influencing poly(A) tail length, we tested for a genetic interaction between*Tdrd5l* and *twin*, which encodes the CCR4 homolog and major deadenylase in *Drosophila* (36). Expression of a weak RNAi trigger for *twin* in the germline of wild-type (wt) females (*nos-Gal4*, UAS-*twin* RNAi) caused no morphological or fertility defects (Fig 3A). Animals heterozygous for a deletion allele of *Tdrd5l* also exhibit no morphological or fertility defects on their own. However, when the *twin* RNAi trigger was expressed in the germline of females heterozygous for a mutation in *Tdrd5l* (*nos-Gal4*, UAS-*twin* RNAi, *Tdrd5l/+*), we observed severely disorganized ovaries that lacked recognizable egg chambers in 100% of ovaries (Fig 3B, N= 20), and these animals were completely sterile. Additionally, to test whether Tdrd5l functions in regulating mRNA stability through removal of the mRNA 5’ cap, we tested for a genetic interaction with *Dcp1*, which encodes one of the major mRNA decapping enzymes in Drosophila (13). As in the case of the *twin* knockdown, there were no morphological or fertility defects when our *Dcp1* RNAi line was expressed in the germline of wt females (*nos-Gal4, UAS-Dcp1 RNAi*, Fig 3C). However, when *Dcp1* was knocked down in the germline of females heterozygous for a mutation in *Tdrd5l*, the ovaries exhibited a severe morphological defect and lacked germline in 100% of ovaries (Fig 3D, N = 20), and these animals were also completely sterile. These dramatic, dose-sensitive genetic interactions between *Tdrd5l* and *twin* or *Dcp1* indicate that Tdrd5l could function in post-transcriptional repression of mRNAs through both the deadenylation and decapping pathways.

**Figure 3:**
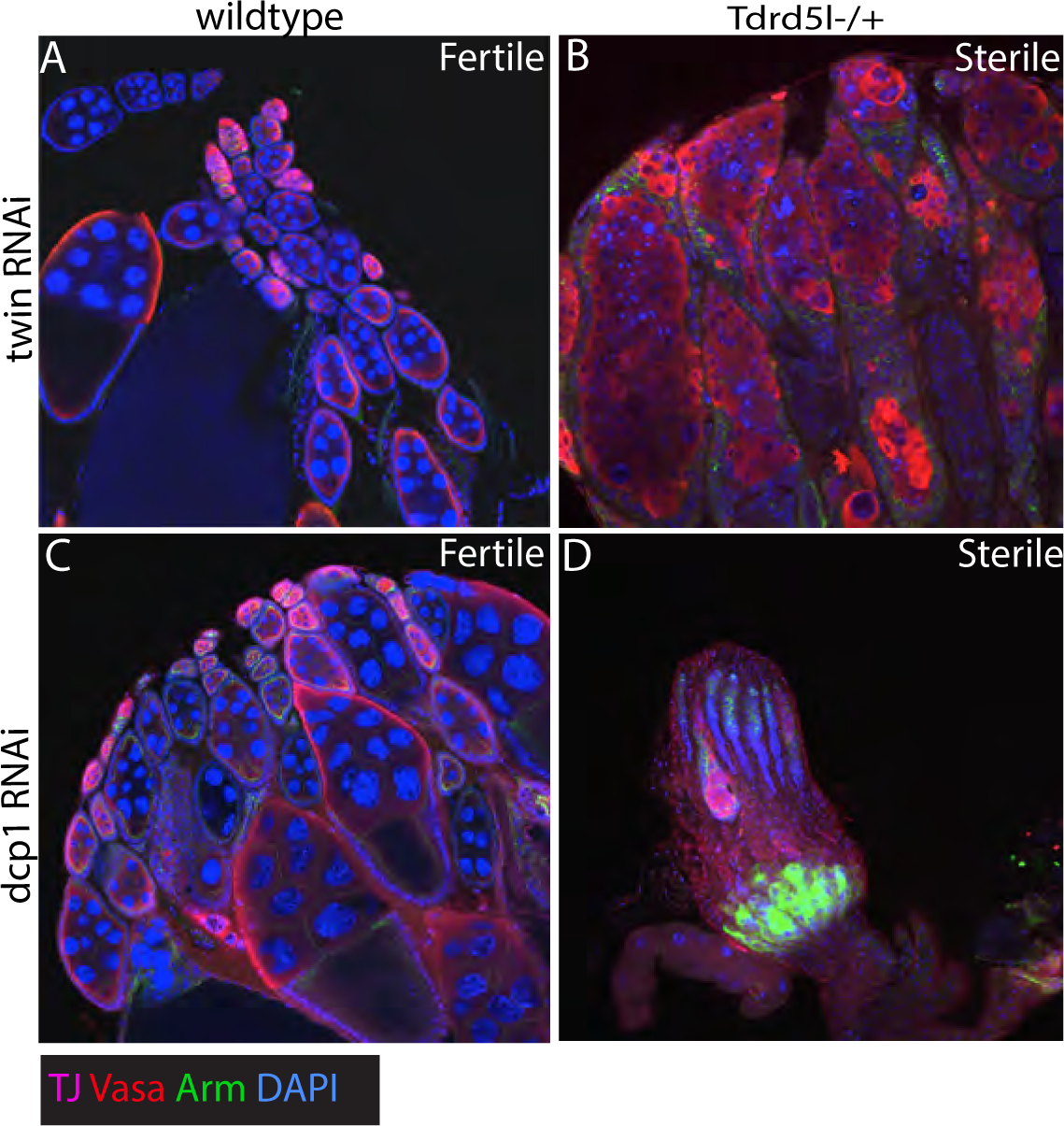
*Tdrd5l* genetically interacts with post-transcriptional gene regulatory factors. (A-D) Immunofluorescence of adult ovaries with genotypes and antibodies as indicated in figure. Fertility status of flies is marked in the top right corner of each panel. (A,B)*nos>twin* RNAi in the germline of wildtype (A) or *Tdrd5l^ΔQ5/+^* heterozygous (B) females. *nos>twin* females (A) have normal ovaries and are fertile while germline development and egg chamber production is severely compromised in *nos>twin, Tdrd5l^ΔQ5/+^* females (B) and these animals are sterile. (C,D) *nos>Dcp1* RNAi in the germline of wildtype (C) or *Tdrd5l^ΔQ5/+^*heterozygous (D) females. *nos>dcp1* RNAi females have normal ovaries and are fertile while germ cells are almost completely absent from *nos>Dcp1*, *Tdrd5l^ΔQ5/+^*females (D) and these animals are sterile.

### Tdrd5l is important for proper egg development

One place where post-transcriptional gene regulation is particularly important is in the developing egg chambers of the female germline. Females trans-heterozygous for predicted null mutant alleles of *Tdrd5l* exhibit a relatively normal ovary morphology but lay reduced numbers of eggs (Fig S6A and Fig 6C). To determine whether the levels of any mRNAs are regulated by *Tdrd5l* we conducted RNAseq in wildtype vs *Tdrd5l*-mutant ovaries. Interestingly, the mRNAs altered in *Tdrd5l* mutants were enriched for genes that are maternally deposited in the embryo, as defined by mRNAs present in 0-2 hr old embryos (37). While maternal genes represent 23.3% of all Drosophila genes, they represent 30% of genes altered in*Tdrd5l* mutants (Fig S7A-B). Further, when the analysis is restricted to genes with a 2-fold or greater change in mRNA level in *Tdrd5l* mutants, the percentage of maternally-expressed genes rises to 94% (Fig S7C). Additionally, we compared our RNAseq data to previously published data identifying RNAs associated with the BicaucalD (BicD)/Egalitarian (Egl) complex involved in transport of RNAs from nurse cells to the oocyte (38). While mRNAs identified in our RNA-seq analysis represent 10.4% of all genes in the Drosophila genome, they represent 30% of the top 100 mRNAs associated with the BicD/Egl complex, further suggesting a role for Tdrd5l in regulating mRNAs that are deposited maternally in the oocyte during oogenesis (Fig S7D).

In *Drosophila*, maternally deposited RNAs are transcribed in the nurse cells where they are silenced until they get to their proper location in the oocyte and are then translationally activated at specific times (39). To investigate whether Tdrd5l plays a role in this process we tested whether the eggs laid by *Tdrd5l*-mutant females exhibited defects in development when crossed to wildtype males. Indeed, we observed that these eggs were greatly reduced in their ability to hatch and give rise to viable larvae (Fig 4A). To determine if the decrease in hatch rate was due to patterning defects, we examined the dorsal appendages (DA) on the offspring, an indicator of Dorsal/Ventral patterning (40). Offspring from trans-heterozygous *Tdrd5l*-mutant females exhibited a greatly increased frequency of DA defects (Fig 4B), which ranged from alterations of DA length (Fig 4D) to a fusion of DA material into a ring (Fig 4E) or a single fused appendage (Fig 4F). If the defects in dorsal-ventral patterning are the major reason behind the embryonic defects, then we would expect that the majority of unhatched embryos would exhibit DA defects. However, when we examined the DA defects specifically of those embryos that failed to hatch, we did not observe a greater percentage of DA defects than we observed when examining all embryos. This indicates that, while DA and D/V defects are the probable cause of death for many of the embryos, other embryos are likely to die for other reasons. When we attempted to examine the cuticle pattern of unhatched embryos we observed no discernable cuticle pattern, indicating that these embryos die prior to cuticle deposition.

**Figure 4:**
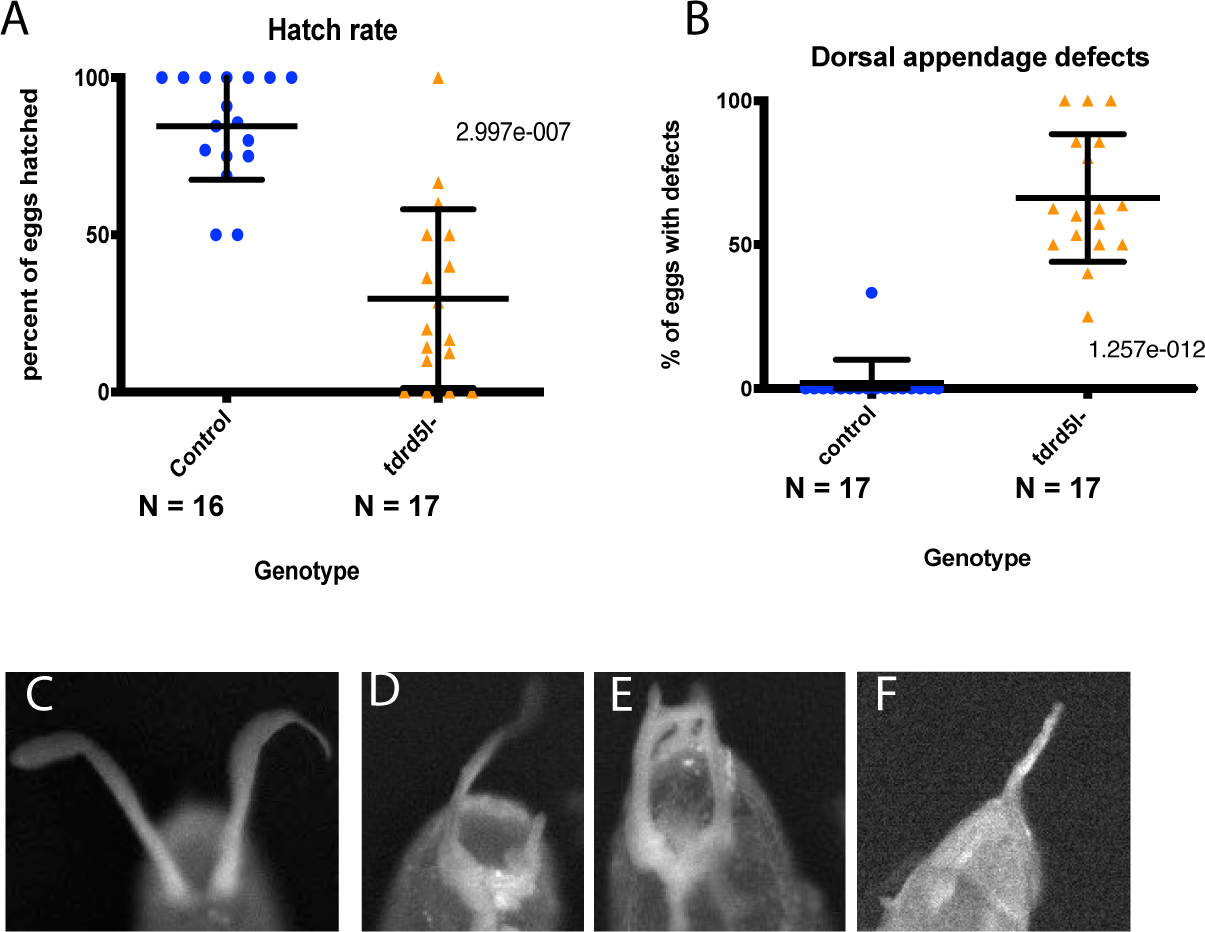
*Tdrd5l* is important for proper egg development. (A) Quantification of hatch rate of eggs laid during single female fecundity assays. Eggs laid by control females (Cas9 ^iso^) are represented by blue dots and eggs laid by *Tdrd5l-*mutant females (Tdrd5l^ΔM4/*Δ*Q5^) are represented by orange dots. B) Quantification of dorsal appendage defects observed in eggs laid during single female fecundity assays. Eggs laid by control females (Cas9^iso^) are represented by blue dots, and eggs laid by *Tdrd5l-*mutant (*Tdrd5l^ΔM4/ΔQ5^*) females are indicated by orange dots. (C-F) Examples of dorsal appendages quantified in B. (C) Example of an egg laid by a *Tdrd5l* mutant female with normal dorsal appendages. (D-F) Examples of eggs laid by *Tdrd5l-* females with dorsal appendage defects.

### *Tdrd5l* is required for repression of Grk expression in nurse cells

The D/V and anterior/posterior (A/P) axes in the developing oocytes are set up by localized translation of maternally deposited RNAs. These RNAs are transcribed in the nurse cells and post-transcriptionally silenced during transport to the oocyte (39). Three of the classic maternally deposited RNAs are *gurken (grk)*, which regulates D/V patterning (20,21), *bicoid (bcd)* which determines anterior fate (24), and *oskar (osk)* which specifies posterior identity and the germ plasm (22). To determine if *Tdrd5l* regulates maternally deposited RNAs we immunostained for the protein products of these three mRNAs in wildtype and *Tdrd5l*-mutant ovaries. Both Grk and Osk protein staining were altered in *Tdrd5l*-mutant ovaries (Fig 5A-H), whereas no Bcd staining was observed in the mutant ovaries, similar to wt (data not shown). In wt ovaries, Grk immunoreactivity is normally observed in the dorsal-anterior corner of the oocyte (Fig 5A, arrowhead) and is absent from the nurse cells of developing egg chambers (Fig 5C). However, in *Tdrd5l* mutant ovaries we observed Grk staining in the nurse cells of mid-stage egg chambers of 80% of ovaries tested (Fig 5D, outlined and inset, N = 25 ovaries), suggesting the *grk* mRNA is no longer translationally repressed in the nurse cells. In addition, we still observed Grk immunoreactivity at the dorsal-anterior corner of developing oocytes (Fig 5B) in *Tdrd5l* mutants.

**Figure 5:**
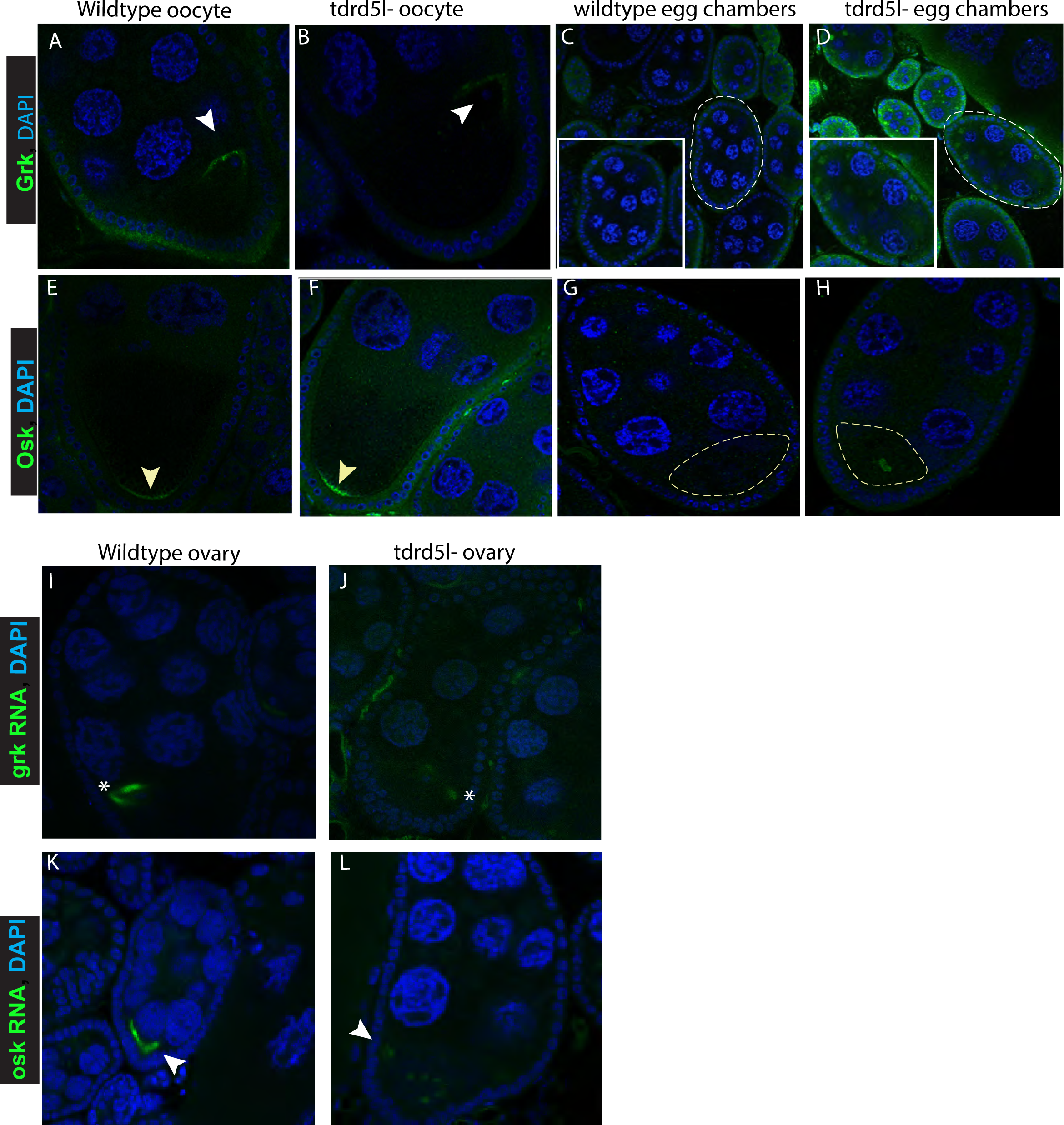
*Tdrd5l* mutants exhibit premature accumulation of maternal proteins. (A-H) Immunofluorescence of proteins produced by maternally deposited RNAs using antibodies as indicated in wildtype (Cas9^iso^) and *Tdrd5l-* (*Tdrd5l^ΔM4/ΔQ5^*) ovaries. (A-D) immunostaining for Grk protein. Grk protein localizes to the dorsal anterior corner of both wildtype (A) and *Tdrd5l-* (B) oocytes in stage 9 egg chambers as marked by the white arrowheads. Grk protein is absent from wildtype nurse cells (C) but is present in*Tdrd5l-*mutant nurse cells (D). Examples of individual Stage 7-8 egg chambers are outlined and enlarged in insets. (E-H) Immunostaining for Osk protein. Osk protein is localized to the posterior of both wild type (E) and *Tdrd5l-* (F) Stage 9-10 oocytes as marked by yellow arrowheads. (G) Osk protein is absent from stage 7 wildtype nurse cells and oocytes. However, Osk protein is observed in the center of 77% stage 7 *Tdrd5l-* oocytes. Oocytes are outlined by dashed yellow line in F and G. (I-L) FISH for maternally deposited RNAs as indicated in wildtype (Cas9^iso^) and *Tdrd5l* mutant (*Tdrd5l^ΔM4/ΔQ5^)* ovaries. (I,J) FISH for *grk* mRNA. *grk* mRNA is localized to the dorsal anterior corner of oocytes in stage 9 egg chambers in both wildtype (I) and*Tdrd5l* mutants (J). The dorsal anterior corner is marked by an asterix. (K,L) FISH for *osk* mRNA. *osk* mRNA is localized to the posterior of wildtype stage 7-8 oocytes (K) but is localized throughout the oocyte in 31% of *Tdrd5l* mutants. Oocytes are marked by white arrowheads

In wt ovaries, Osk immunostaining is normally restricted to the posterior pole of developing oocytes beginning at stage 10 (Fig 5E, arrowhead) (22). In stage 10 *Tdrd5l-*mutant egg chambers we still observed Osk protein correctly localized to the posterior pole (Fig 5F). However, in 77% of stage 9 *Tdrd5l*-mutant egg chambers we observed premature staining for Osk, and in these cases Osk protein was localized to the middle of the oocyte rather than the posterior end (Fig 5H, N = 25 ovaries). However, unlike with Grk, we did not observe Osk protein expressed in the nurse cells of *Tdrd5l* mutants (Fig 5H).

To determine if the ectopic expression of Grk and Osk proteins was due to defects in RNA transport from the nurse cells or RNA localization in the oocyte, we conducted fluorescent *in situ* hybridization (FISH) to visualize the *grk* and *osk* mRNAs (Fig 5I-L). We observed no changes in *grk* mRNA localization between wildtype and *Tdrd5-*mutant ovaries (Fig 5I-J). Contrary to what we observed with the *grk* RNA, we observed altered *osk* mRNA localization in *Tdrd5l*-mutant oocytes (Fig 5L). In *Tdrd5l* mutant oocytes, *osk* mRNA was not tightly localized to the posterior and was instead observed throughout the oocyte in a 31% of ovaries (Fig 5L, N= 15 ovaries).

Grk is first required at the posterior of the oocyte to specify the posterior follicle cells, which are then required for proper microtubule orientation and localization of*osk* RNA and protein to the posterior pole (41). Thus, it is possible that the mis-localization of*osk* we observed could be secondary to defects in *grk* function. To test whether the microtubule network is properly oriented in *Tdrd5l* mutants, we used Kinesin-LacZ (Kin-LacZ) to mark the plus ends of the microtubules (21). In wt ovaries the microtubule network in the oocyte is polarized with plus ends at the posterior, therefore Kin-LacZ should be localized tightly to the posterior by stages 9 and 10 of egg chamber development (Fig S8A white arrow, and Fig S8B yellow arrow, quantified in Fig S8E). In *Tdrd5l*-mutant ovaries we detected a range of Kin-LacZ localization: some oocytes had tight posterior localization, while others had some posterior localization of Kin-LacZ with plumes of staining extending away from the posterior pole. In the most extreme cases we observed diffuse Kin-LacZ localization in the center of the oocyte (Fig S8C white arrow and Fig S8D yellow arrow, quantified in Fig S8E). These data indicate that the defects in *osk* posterior localization may be caused by loss of *grk* function rather than defects in the direct regulation of *osk* itself.

### *Tdrd5l* represses Orb protein expression in nurse cells

Work by other labs has shown that Grk translation in the oocyte is activated by Oo18 RNA binding protein (Orb) (42–44). Orb is a cytoplasmic polyadenylation element binding protein (CPEB) that is highly expressed in the oocyte where it recruits Wispy to lengthen the*grk* poly(A) tail to activate its translation (43). It has also been shown that ectopic expression of Orb in the nurse cells is sufficient to prematurely activate Grk translation (44). To test whether ectopic expression of Orb in the nurse cells could be the cause of Grk translation in*Tdrd5l*-mutant nurse cells, we stained for Orb protein in *Tdrd5l*-mutant ovaries. As previously described, in all wt ovaries we observed Orb immunofluorescence primarily in the oocyte and it was largely absent from the nurse cells of later egg chambers, except the nurse cell closest to the oocyte (Fig 6A, squared region). In 100 % of *Tdrd5l*-mutant ovaries we saw an expansion of Orb protein expression into almost all the nurse cells suggesting Orb translation had been de-repressed in *Tdrd5l* mutants (Fig 6B, squared region, N= 26).

**Figure 6:**
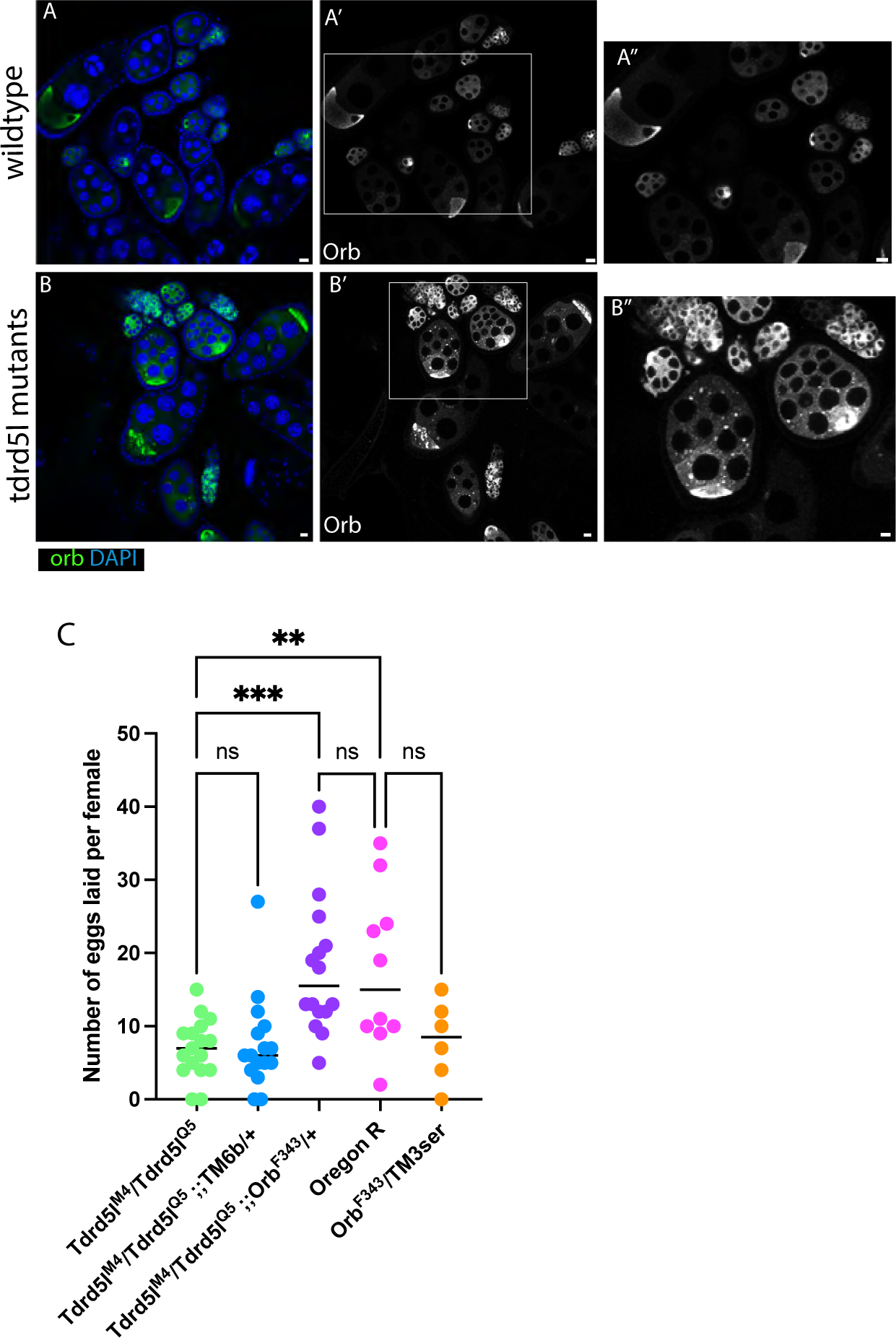
Tdrd5l represses Orb translation. (A-B) Immunofluorescence of Orb in wildtype (Cas9^iso^) and *Tdrd5l* mutant (*Tdrd5l^ΔM4/ΔQ5^*) ovaries. Scale bars = 5 microns. A) Orb protein expression is detected at high levels in wildtype oocytes. (A’) boxed area is enlarged in (A’’). Orb protein expression is detected at high levels in both oocytes and nurse cells of *Tdrd5l-* ovaries(B). B’) boxed area is enlarged in (B’’). C) Quantification of egg laid by females from an *Orb/Tdrd5l* genetic interaction assay. Fewer eggs are laid by *Tdrd5l-* (*Tdrd5l^ΔM4/ΔQ5^*) females than by *Tdrd5l-* mutant females transheterozygous for an *orb*^F343^ allele (*Tdrd5l^ΔM4/ΔQ5^;; Orb^F343/+^*). Note that eggs laid by transheterozygous flies are at a similar level to eggs laid by wildtype (OregonR) flies. P-value ≤ 0.01 is denoted by **. P-value ≤ 0.001 is denoted by ***.

To test whether the phenotype observed in *Tdrd5l* mutants is due to increased Orb expression, we determined whether reduction of *orb* function could suppress the *Tdrd5l*-mutant phenotype. *Tdrd5l*-mutant females that were also heterozygous for an *orb* loss of function allele (*Tdrd5l-/Tdrd5l-; orbF343/+* females) laid significantly more eggs than *Tdrd5l-* mutant females, and a similar number of eggs to wt control females (Fig 6C). However, being heterozygous for an *orb* mutant allele did not rescue the egg hatching defect observed in *Tdrd5l-*mutants (not shown). This suggests that the defects observed in *Tdrd5l* mutants are partially due to the increased Orb expression observed in the nurse cells, but other defects may be due to misregulation of other RNAs or other functions of Tdrd5l.

## Discussion

Work from a number of labs has demonstrated the importance of RNA granules for proper germ cell identity and function. Here we have extended our study of Tdrd5l and shown that it localizes to the periphery of a novel germline structure, the “Tdrd5l body”, that does not exhibit the characteristics of any previously identified germline granule. While we previously demonstrated that Tdrd5l expression is highly male-biased in the undifferentiated germline, we show here that Tdrd5l bodies are also present during germline differentiation in females.*Tdrd5l* does not regulate transposon expression, like other Tdrd5 family members, but instead regulates the expression of maternal RNAs that need to be silenced in nurse cells as they are transported to the oocyte.

### Analysis of the “Tdrd5l body”

Previous work studying expression of the Tdrd5l protein used a genomic BAC with an N-terminal epitope tag that we have subsequently found interferes with*Tdrd5l* function. Therefore, we used an internal epitope-tag placed into the endogenous *Tdrd5l* locus that retains full *Tdrd5l* function, as well as a peptide polyclonal antibody generated against native Tdrd5l, to characterize the localization of Tdrd5l. We were able to verify that Tdrd5l is more highly expressed overall in the male germline than the female germline, and is expressed in male GSCs but is not observed in female GSCs (Fig 1). This is consistent with *Sex lethal* being a repressor of Tdrd5l expression as we previously found (25). However, we also describe the expression of Tdrd5l in the female germline, where it is found in germ cells that have begun to differentiate in the germarium (Fig 1C-D), and in nurse cells and oocytes in the developing egg chambers (Fig 1E-F). Interestingly, in both male and female germ cells, the punctae of Tdrd5l expression appear “hollow” indicating that Tdrd5l resides on the surface of the structures it identifies.

Our analysis also indicates that the Tdrd5l bodies are not representative of any type of previously described germline granule in *Drosophila*. The most prominent germline granule is the perinuclear nuage which contains the helicase Vasa and other factors important for piRNA biogenesis, like Aub and Ago3. While the Tdrd5l bodies are often observed in a perinuclear location, they notably lack Vasa and are not enriched for Aub or Ago3 (Fig 2). Thus, while Tdrd5l bodies may interact with the nuage, they appear to be distinct entities. A large granule, the piRNA nuage giant body, has been described that is associated with the nuage (45). However, unlike the Tdrd5l body, this body contains Vasa and appears only in primary spermatocytes, and so is distinct from the Tdrd5l body. The localization of the Tdrd5l body is also not consistent with it being part of the germ plasm or sponge bodies found in the oocyte (46–49). The Tdrd5l body appears to be distinct from RNA granules found more commonly in different cell types, such as P bodies, U bodies and stress granules. P bodies characteristically contain Dcp1 and are much smaller in size than Tdrd5l bodies. Our previous work suggested some overlap in localization between Tdrd5l and Dcp1 (25) but the more detailed analysis described here reveals that smaller, Dcp1-positive bodies can be seen associating with Tdrd5l bodies, but that Dcp1 is not present in the Tdrd5l bodies themselves (Fig 2). We also do not see overlap between Tdrd5l and the U body protein Smn (Fig S4). Lastly, the Tdrd5l bodies are observed constitutively in the germline and are not affected by stresses such as starvation or changes in temperature (C.P. unpublished data) indicating that they are not stress granules.

Germline granules have been most well-studied in C. elegans, and a number of distinct regions of the nuage have been described in this species (50,51). At many stages of germline development, P granules are associated with the nuclear periphery and, like Drosophila nuage, contain Vasa-class helicases (the GLH’s) as well as Ago proteins (52–54). However, associated with the P granules are other perinuclear regions known as the Mutator foci, SIMR foci and the Z granules, that have distinct protein components (51). One possibility is that the Tdrd5l body represents a similar sub-structure associated with the nuage in *Drosophila*. In addition to studying the function of *Tdrd5l*, identifying other proteins and possibly RNAs that are present in the Tdrd5l body will be an important next step.

### Function of the Tdrd5 proteins

There are many types of Tudor domain containing proteins and they can be classified into sub-families based on the homology of their Tudor domains. In diverse animal species, including mouse and humans, there is a single Tdrd5 protein. Interestingly, flies have two such proteins, Tdrd5l and Tejas (10,25). Many Tdrd5 proteins, like mouse TDRD5 and *Drosophila* Tejas, also contain an N-terminal LOTUS domain which is known to bind Vasa-type helicases (27), but Drosophila Tdrd5l lacks this domain. However, the N-terminus of Tdrd5l must be important for its function since placing even a small epitope tag in this position compromises Tdrd5l function. Consistent with its Vasa-binding LOTUS domain, Tejas associates with the Vasa-positive nuage where it acts in the piRNA pathway to repress germline transposon expression (10). In contrast, we did not observe Tdrd5l co-localizing with the nuage (Fig 2A), and we have observed no change in transposon expression in *Tdrd5l* mutants (Fig S5). Instead, we find that *Tdrd5l* exhibits potent genetic interaction with pathways that repress mRNAs at the post-transcriptional level, including the mRNA decapping and deadenylation pathways (Fig 3). This is also consistent with a role for Tdrd5l in regulating mRNA repression in nurse cells (see below). Thus, it may be that Tejas is more specific for transposon regulation while Tdrd5l is involved in post-transcriptional regulation of germline mRNAs.

Mouse TDRD5 associates with the Chromatoid Body, a nuage-related structure. In addition, mice mutant for *Tdrd5* exhibit defects in transposon regulation and spermatogenesis (26), similar to Drosophila *tejas* (10). Interestingly, *Tdrd5*-mutant mice also exhibit defects in regulation of pachytene piRNAs (55,56), which are proposed to regulate mRNAs important for spermatogenesis rather than to repress transposons. One intriguing hypothesis is that the role of the single TDRD5 protein in mice in regulating both transposon repression and mRNA expression has been divided between the two Tdrd5 proteins in Drosophila, Tejas and Tdrd5l. In addition, both Drosophila proteins have functions in oogenesis in addition to spermatogenesis (10,25, this work). Similarly, knock down of a Tdrd5 homolog in Locust also affects both oogenesis and spermatogenesis (57). While a role for mouse TDRD5 in the female germline has not been described, consortium data indicate that *Tdrd5* is expressed in the mouse ovary (58), making this an interesting place to look for additional functions of TDRD5 in the mouse.

### Regulation of Maternally deposited RNAs

Prior work from our lab demonstrated that *Tdrd5l* promotes male identity in the germline. Consistent with this, Tdrd5l is expressed in male GSCs but is repressed in female GSCs by the action of *Sxl* (25). However, Tdrd5l is also expressed in differentiating germ cells in both males and females ((25), and Fig 1). This suggests that Tdrd5l might promote male identity in early germ cells and germline stem cells but also regulate aspects of germline differentiation in both sexes. One process in the female germline that relies heavily on post-transcriptional regulation of RNAs is the production and transport of maternal RNAs from nurse cells into the oocyte.

Maternal contribution of RNAs, proteins, and organelles is conserved from flies through vertebrates (59,60), and supports embryonic patterning and embryogenesis prior to activation of the zygotic genome and beyond. In *Drosophila*, maternal RNAs are transcribed in the nurse cells, and are post-transcriptionally silenced to prevent their translation during transport to the oocyte and prior to the specific time they should be activated. The cytoplasmic polyadenylation element binding protein (CPEB) Orb is known to activate translation of maternal RNAs such as *osk* and *grk* in the oocyte (43,44,61). Like other maternal RNAs, *orb* is transcribed in nurse cells and transported to the oocyte where it is translated. Interestingly, Orb/CPEB homologs are critical for regulating translation of maternal RNAs in vertebrate systems as well (62).

We observed defects in expression of Orb, Grk and Osk proteins in *Tdrd5l* mutants (Fig 5 and Fig 6). We saw ectopic accumulation of both Orb and Grk in nurse cells (Fig 5D and Fig 6B), while Osk protein was sometimes observed in the middle of the oocyte instead of its normal location at the posterior pole (Fig 5H). We also observed maternal-effect lethality of embryos from *Tdrd5l*-mutant mothers that included dorsal appendage defects consistent with Grk’s role in dorsal-ventral patterning. One possible explanation for all of these phenotypes is a failure of Orb to be properly repressed in nurse cells. Ectopic Orb translation in nurse cells can cause improper activation of Grk translation in these cells (44), and this might also lead to insufficient translation of Grk in the oocyte. Early Grk expression at the posterior pole of the oocyte is necessary for proper posterior determination and defects in this process can lead to disruption of *osk* mRNA and protein localization to the posterior pole (19,21) as observed in *Tdrd5l* mutants (Fig 5L). Further, impaired Grk expression in the dorsal-anterior corner of the oocyte can cause problems in specification of dorsal follicle cells, and defects in dorsal appendage formation and dorsal-ventral patterning of the embryo. Thus, one primary defect in*Tdrd5l* mutants could be a failure to properly repress *orb* translation in the nurse cells. In agreement with this possibility we see a partial rescue of the *Tdrd5l* egg-laying defect when *orb* function is compromised (Fig 6).

Other factors have been implicated in repressing *orb* translation in nurse cells, including FMR1, the Drosophila homolog of Fragile X mental retardation protein (63), and Cup, a translational repressor (64). Mutations in both *FMR1* and *cup* cause Orb accumulation in nurse cells similar to what we observe in *Tdrd5l* mutants. One intriguing hypothesis is that the Tdrd5l body in nurse cells is a site where mRNAs such as *orb* get marked for translational repression by factors such as FMR and Cup. Since the nuage and nuage-related material in flies, worms and mice all function in small RNA regulatory pathways, the marking of mRNAs like*orb* in the Tdrd5l granule may also involve such regulatory RNAs.

## Materials and Methods

### Fly stocks and CRISPR tagging

Fly stocks used in this paper were obtained from the Bloomington stock center unless otherwise noted. *Nos*-gal4 (65), *twin* RNAi(BDSC# 32901), *Dcp1* RNAi(BDSC# 67874), *mcherry* RNAi(BDSC# 35785), *orb* RNAi(BDSC# 43143), Vasa-Cas9(BDSC# 56552), piggyBac (BDSC# 32070), Kin-LacZ (T. Schupbach). *Tdrd5l* mutant alleles were generated previously by our lab (25). *Tdrd5l*:GFP and *Tdrd5l*:FLAG alleles were generated using CRISPR as described by the fly CRISPR group (66). The GFP sequence followed by a linker sequence was inserted in place of the Tdrd5l start codon. The internal flag construct contains a 3x FLAG sequence flanked by linkers in the middle of exon 3 following K225. The GFP and FLAG donor plasmids and pU6 gRNA plasmid were obtained from the *Drosophila* genetics resources center and were stock numbers 1365, 1367, 1363 respectively.

### Immunofluorescence and antibody generation

All gonads were dissected, and fixed, and stained as previously published (67). All images were taken on a Zeiss LSM 700, LSM 800 with airyscan detector (when noted), or LSM 980 with airyscan detector (when noted). Primary antibodies used were Guinea pig anti-Tdrd5l 1:10,000 (this paper), rat anti-HA 1:100 (Roche Cat # 3F10), rabbit anti-Vasa 1:10,000 (R, Lehman), Guinea pig anti-TJ 1:1000 (M. Van Doren), mouse anti Armadillo 1:100 (DSHB, N2 7A1), Mouse anti Orb 1:40 (DSHB, 6H4), mouse anti-Gurken 1:20 (DSHB, 1D12), rabbit anti-oskar 1:1000 (A. Ephrussi), rat anti-Ago3 1:500 (T. Kai), mouse anti-Aub 1:1000 (MC. Siomi). Secondary antibodies were used at 1:500 (Goat anti-rabbit alexa-fluor 546 Cat # A11011, Goat anti-rabbit alexa-fluor 488 Cat # A11034, Goat anti-mouse alexa-fluor 488 Cat # A11029, Goat anti-mouse alexa-fluor 546 Cat # A11004, Goat anti-mouse alexa-fluor 633 Cat # A21050, Goat anti-rat alexa-fluor 488 Cat # A11006. Goat anti-rat alexa-fluor 633 Cat # A21094, Goat anti-guinea pig alexa-fluor 488 Cat # A11073, Goat anti-guinea pig alexa-fluor 633 Cat # A21105). Samples were stained in DAPI solution and mounted in DABCO.

### Quantification of nuage association

To quantify the association of Tdrd5l granules with Vasa, we calculated the number of Tdrd5l granules in the 8 and 16 cell cysts that localized either to the nuclear periphery (perinuclear) or to the cytoplasm. For each group we then calculated the number of Tdrd5l granules that are directly adjacent to Vasa granules, co-localize with Vasa, or do not associate with Vasa at all.

### Egg lay assays

Trans-heterozygous *Tdrd5l* mutant females and control females were aged 7days with *OregonR* males. Individual females and males were placed in condos on grape juice plates with wet yeast paste for 24hrs, and then taken off the plate. The number of eggs laid and those with dorsal appendage defects were counted for each female. At 48hrs these counts were done again to count how many had hatched.

### Genetic interaction assays

UAS-RNAi lines were crossed to Nos-gal4 for controls or *Tdrd5l^Q5^;;nos-gal4* for mutants. *twin* and *Dcp1* were knocked down using a UASp-shRNAi driven in the germline by nos-gal4 in a wildtype background and in a *Tdrd5l-* background. female progeny were heterozygous (*Tdrd5l/+*), Progeny were aged 5-10days then dissected immunostained as described above using anti-vasa, anti-TJ, anti-Arm and DAPI.

### Fluorescence in situ hybridization

FISH for *gurken* and *oskar* were conducted as published by the Berg lab (68). cDNA clones 2169 and 7305 for probes were obtained from the Drosophila Genetics Resource Center.

### RNA sequencing

For testis RNA sequencing, Testes were dissected from 5d old males, RNA was prepped using RNA-Bee(Tel-test) 3 biological replicated were used for each genotype. Libraries were prepped using a previously established protocol (69). 100bp paired end sequencing was conducted by the Johns Hopkins Genetics Resources Core facility For ovary RNA sequencing, ovaries were dissected from virgin females, and 3 biological replicates were dissected per genotype. RNA was prepped using The Direct-zol RNA micro prep kit (Zymogen). Library construction and 100bp paired end sequencing was conducted by the Johns Hopkins Genetics Resources Core facility.

Read quality for both the ovary and testis RNAseq data was determined using the fastQC kit. Reads were mapped using Tophat and HTSeq to the *Drosophila* genome using Ensemble release BDGP6. Differential gene expression analysis was conducted using DESeq.

Differential transposon analysis was carried out as previously (70). Briefly, RNAseq reads were mapped to the FlyBase r6.46 genome using Hisat2 (-k 1--rna-strandness RF--dta) (PMID:35266522, PMID: 25751142). Read counts for transposons were determined based on their genomic locations according to the UCSC RepeatMasker track and read counts were analyzed with HTSeq-count (PMID: 29106570, PMID: 25260700). Differential expression analysis of transposons was determined with DESeq2 (pAdjustMethod = ‘fdr’)(PMID: 25516281).

Strains and plasmids are available upon request. The authors affirm that all data necessary for confirming the conclusions of the article are present within the article, figures, and tables.

## Acknowledgements

We thank the *Drosophila* community, the Bloomington stock center, DRGC, and Developmental Studies Hybridoma Bank for stocks and reagents. We would also like to thank the Chou lab for Dcp1-YFP flies, as well as Trudi Schupbach and Liz Gavis for reagents and helpful discussion on this work. We thank members of the Johns Hopkins community; the Johnston lab, and John Kim for helpful discussion as well as Fred Tan for help with bioinformatics. This work was supported by NIH grant R21HD101781 to MVD.

---

**Supplemental Figure 1:**
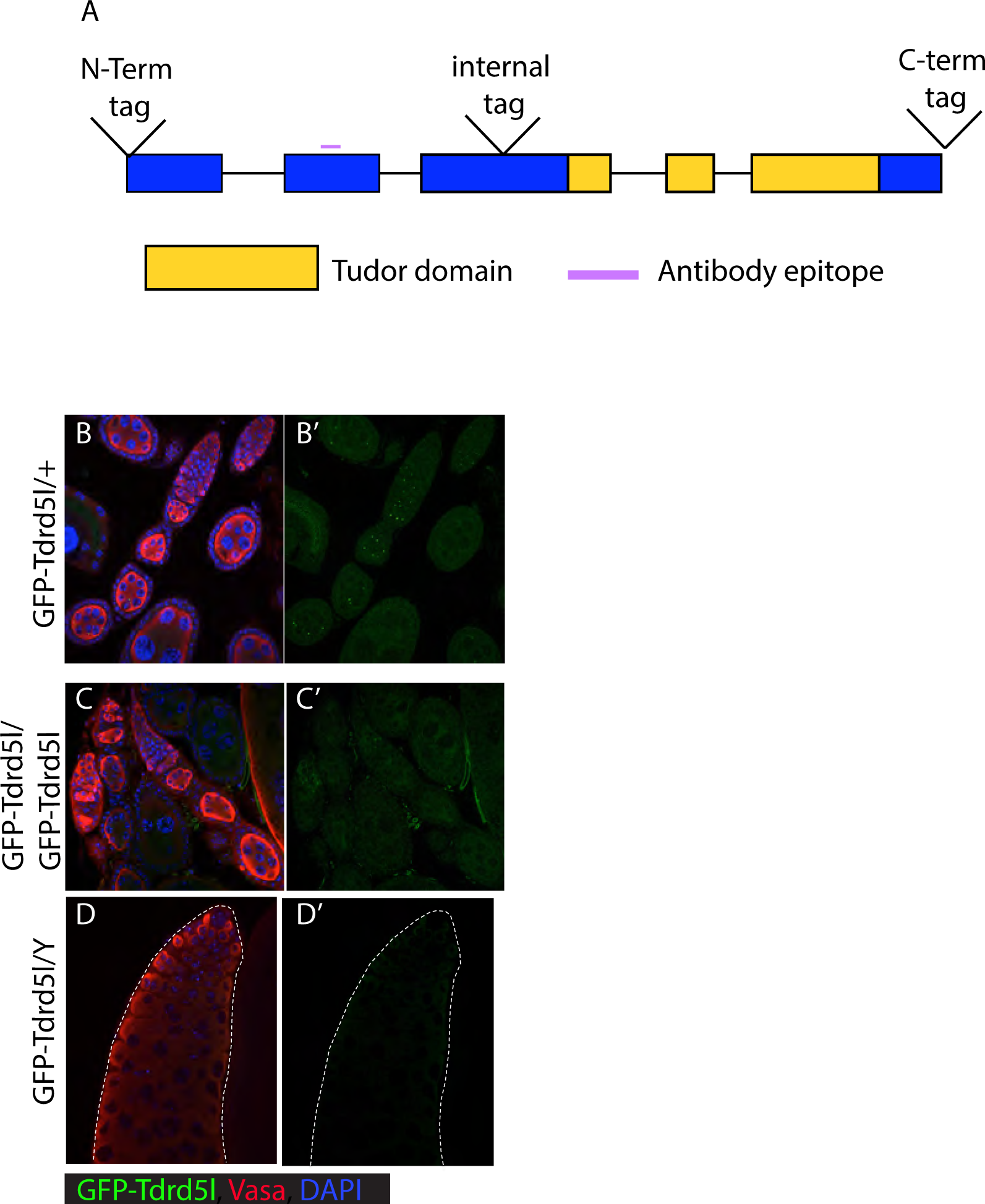
Tdrd5l localization to cytoplasmic bodies in the *Drosophila* germline is dependent on the N-terminus. (A) Schematic of the *Tdrd5l* gene locus, insertion sites for endogenous tags are marked as N-Term tag, Internal tag, and C-Term tag. Antibody was raised against the epitope marked by the purple line. (B-D) Immunofluorescence of adult gonads expressing endogenous Tdrd5l N-terminally tagged by GFP. Antibodies used as indicated. (B) Tdrd5l-GFP expression in ovaries heterozygous for *GFP-Tdrd5l/untagged-Tdrd5l*. Note the localization of GFP-Tdrd5l to cytoplasmic bodies. (C) GFP expression in ovaries homozygous for GFP-Tdrd5l. Note the lack of GFP signal. (D) GFP expression in males hemizygous for GFP-Tdrd5l (*GFP-Tdrd5l/Y*). Note the lack of GFP signal. The testis is outlined by a white dotted line in (D’).

**Supplemental Figure 2:**
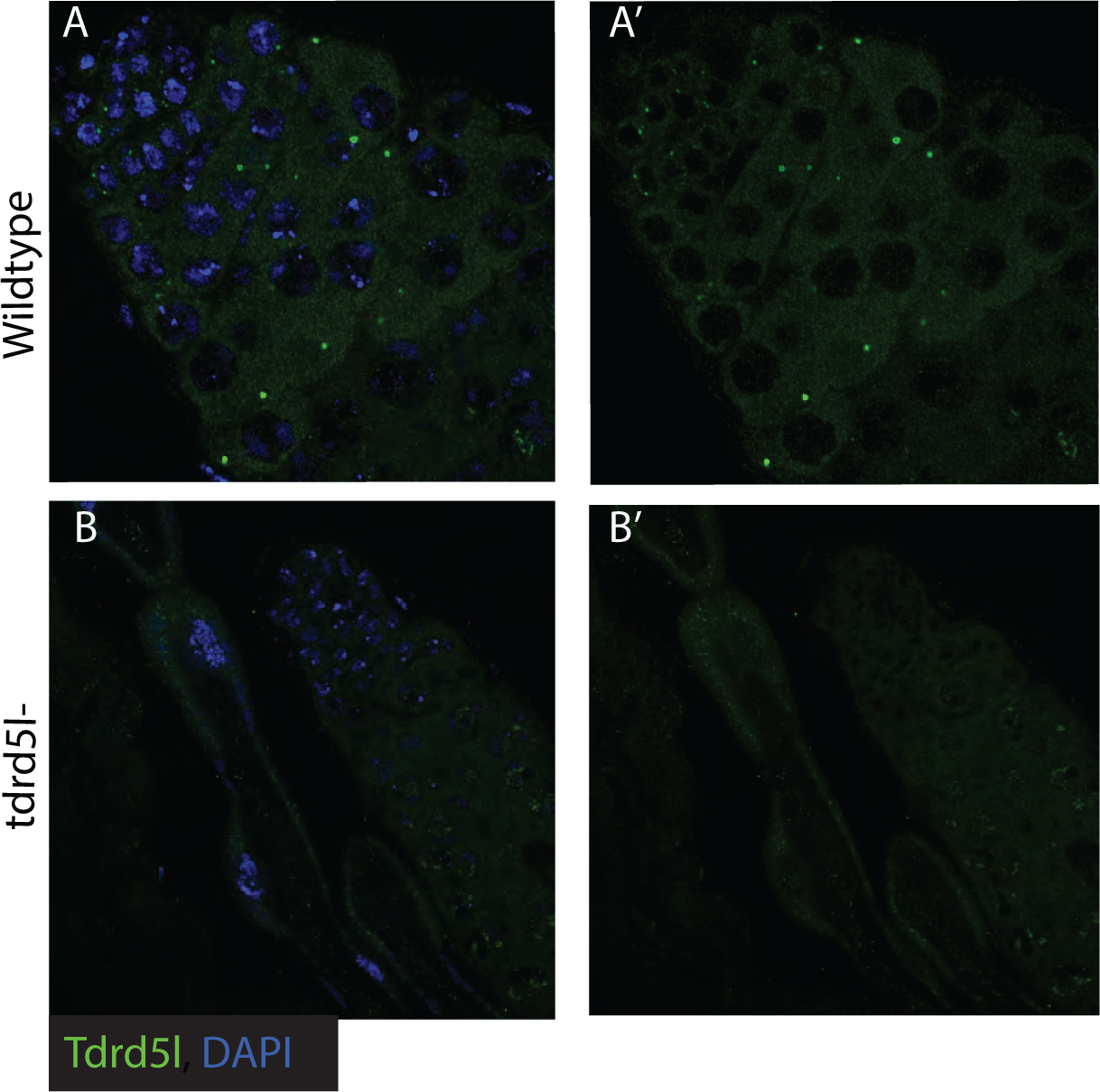
Validation of Tdrd5l antibody. (A-B) Immunofluorescence of the anti-Tdrd5l antibody in adult testes using antibodies as indicated. Tdrd5l expression is detected in wildtype testes (Oregon R) where the antibody recognizes Tdrd5l bodies (A). Tdrd5l expression was not detected in *Tdrd5l-* (*Tdrd5l^Δ^*^M4^) testes (B).

**Supplemental Figure 3:**
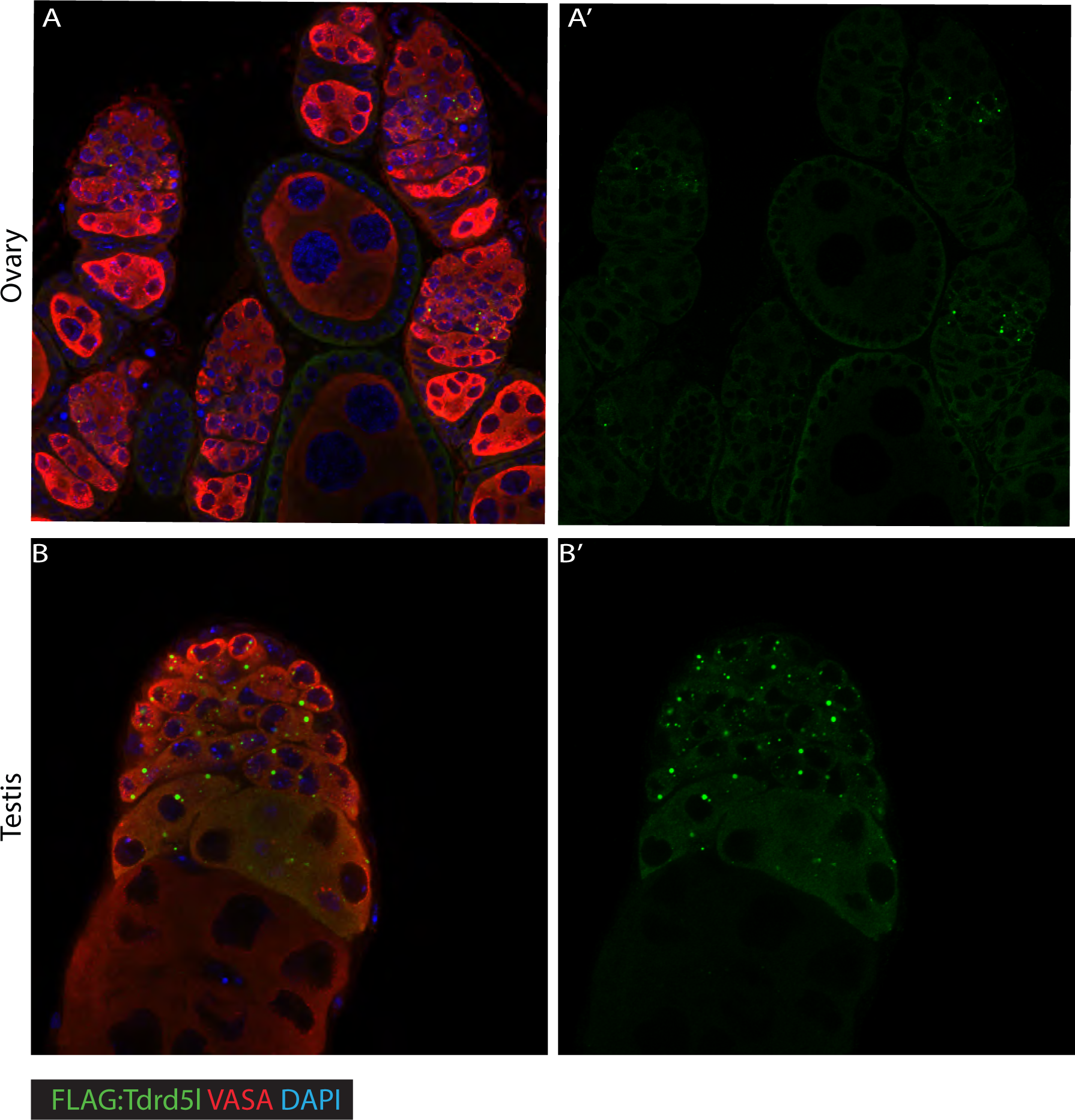
Tdrd5l expression is higher in males than females. A-B) Immunofluorescence of internal FLAG-Tdrd5l using the same confocal settings to image male and female gonads. FLAG-Tdrd5l is expressed at higher levels in testes (A) compared to Tdrd5l-FLAG staining in ovaries (B) when imaged with the same confocal settings. Note the localization of Tdrd5l to cytoplasmic bodies in both ovaries and testes.

**Supplemental Figure 4:**
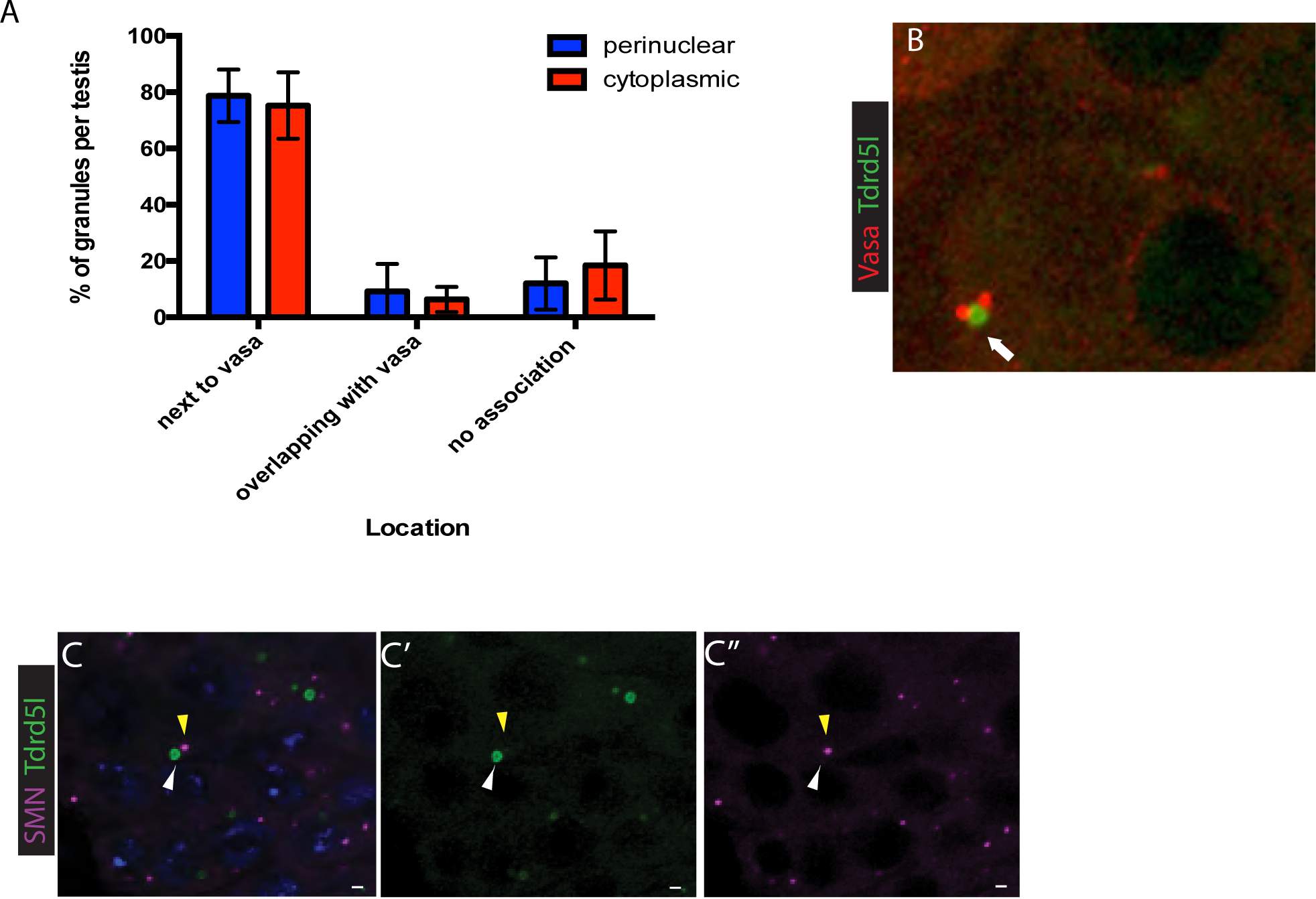
Tdrd5l localizes to an unknown body. (A) Quantification of Tdrd5l body association with Vasa stained nuage. Red bars represent cytoplasmic Tdrd5l bodies while blue bars represent perinuclear Tdrd5l bodies. Bodies were classified as localizing adjacent to Vasa as in (B), overlapping with Vasa, or having no association with Vasa. (B) Example of a cytoplasmic Tdrd5l body visualized using an internal FLAG tagged allele of Tdrd5l localized next to Vasa-positive granules. (C) Immunofluorescence of FLAG-Tdrd5l and HA-SMN. Tdrd5l bodies shown in green and marked by a white arrowhead. SMN positive U-bodies are shown in magenta and marked by a yellow arrowhead. Note Tdrd5l bodies do not co-localize with SMN positive U-bodies.

**Supplemental Figure 5:**
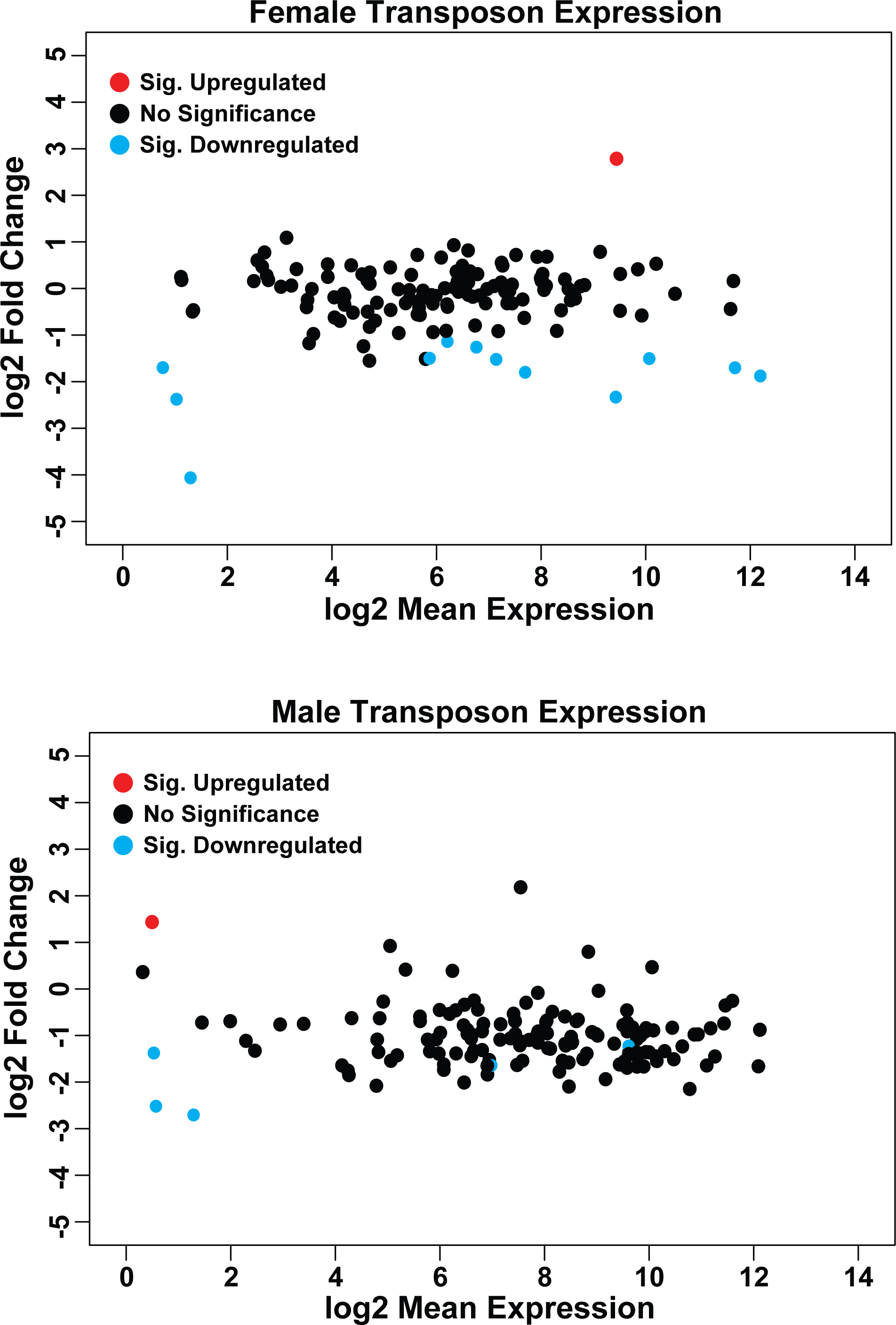
Tdrd5l does not repress transposon expression. (A-B) Differential expression analysis of transposons in *Tdrd5l-* gonads (Females = *Tdrd5l^ΔM4/ΔQ5^*, Males = *Tdrd5l^ΔM4^/Y*) compared to wt. Each dot signifies a single transposon gene. Blue dotes are down-regulated genes, red dotes are upregulated-genes, and black dots are transposons with no change in expression levels. A) RNA sequencing of *Tdrd5l* mutant ovaries compared to wildtype (Cas9^iso^) ovaries showed no de-repression of transposon expression. B) RNA sequencing of *Tdrd5l* mutant testes compared to wildtype (Cas9^iso^) testes showed no global changes in transposon expression.

**Supplemental Figure 6:**
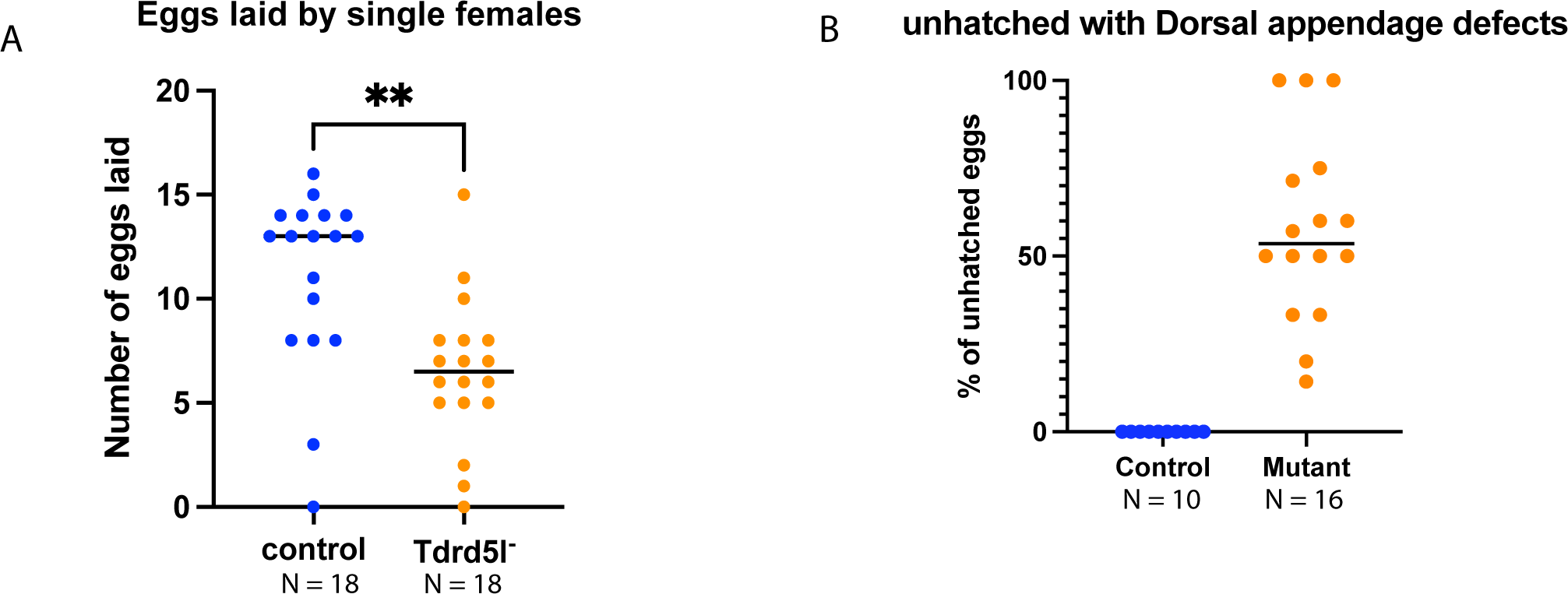
Tdrd5l is important for proper egg development. (A-B) Quantification of fecundity assays performed using single female egg lay assays. (A) Eggs laid per *Tdrd5l-* (*Tdrd5l^ΔM4/ΔQ5^*) female (orange dots) compared to control (Cas9^iso^) females (blue dots). Note the decreased number of eggs laid by *Tdrd5l*-mutants. (B) Number of unhatched embryos with dorsal appendage defects per *Tdrd5l-* (*Tdrd5l^ΔM4/ΔQ5^*) female (orange dots) compared to wt control (Cas9^iso^, blue dots). Note that the % of unhatched embryos with Dorsal Appendage defects is similar to the % of ALL embryos with Dorsal Appendage defects (Fig 4B), indicating that other defects in addition to D/V patterning defects affect viability of embryos from *Tdrd5l-*mutant mothers.

**Supplemental figure 7:**
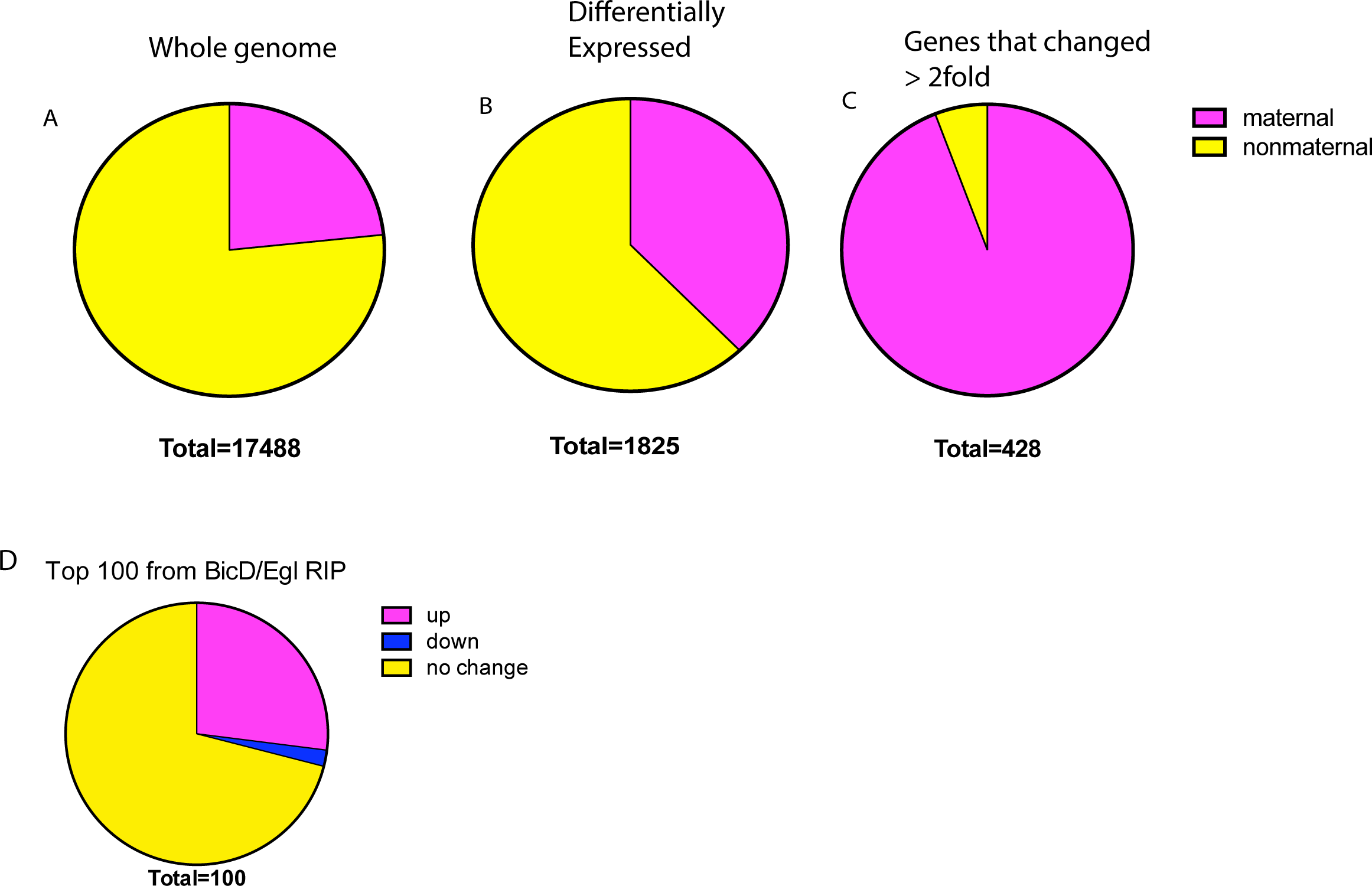
Tdrd5l regulates maternally deposited RNAs. (A-C) Quantification of changes in maternal RNA expression in *Tdrd5l-* (*Tdrd5l^ΔM4/ΔQ5^*) ovaries compared to control (Cas9^iso^) ovaries. Maternally deposited RNAs are defined as RNAs found in 0-2hr embryos from modENCODE. In each pie chart the share of maternal RNAs is shown in magenta, while the share of non-maternal RNAs is shown in yellow. (A) 23% of all genes in the genome are maternally deposited. (B) 30% of genes differentially expressed in *Tdrd5l-*mutant (*Tdrd5l^ΔM4/ΔQ5^*) ovaries are maternally deposited. (C) 90% of genes with a greater than 2-fold change in gene expression in *Tdrd5l*-mutant (*Tdrd5l^ΔM4/ΔQ5^*) ovaries are maternally deposited. (D) 25% of RNAs pulled down in a previously published BicD/Egl RIP-seq data set are upregulated in *Tdrd5l-*mutant (*Tdrd5l^ΔM4/ΔQ5^*) ovaries.

**Supplemental figure 8:**
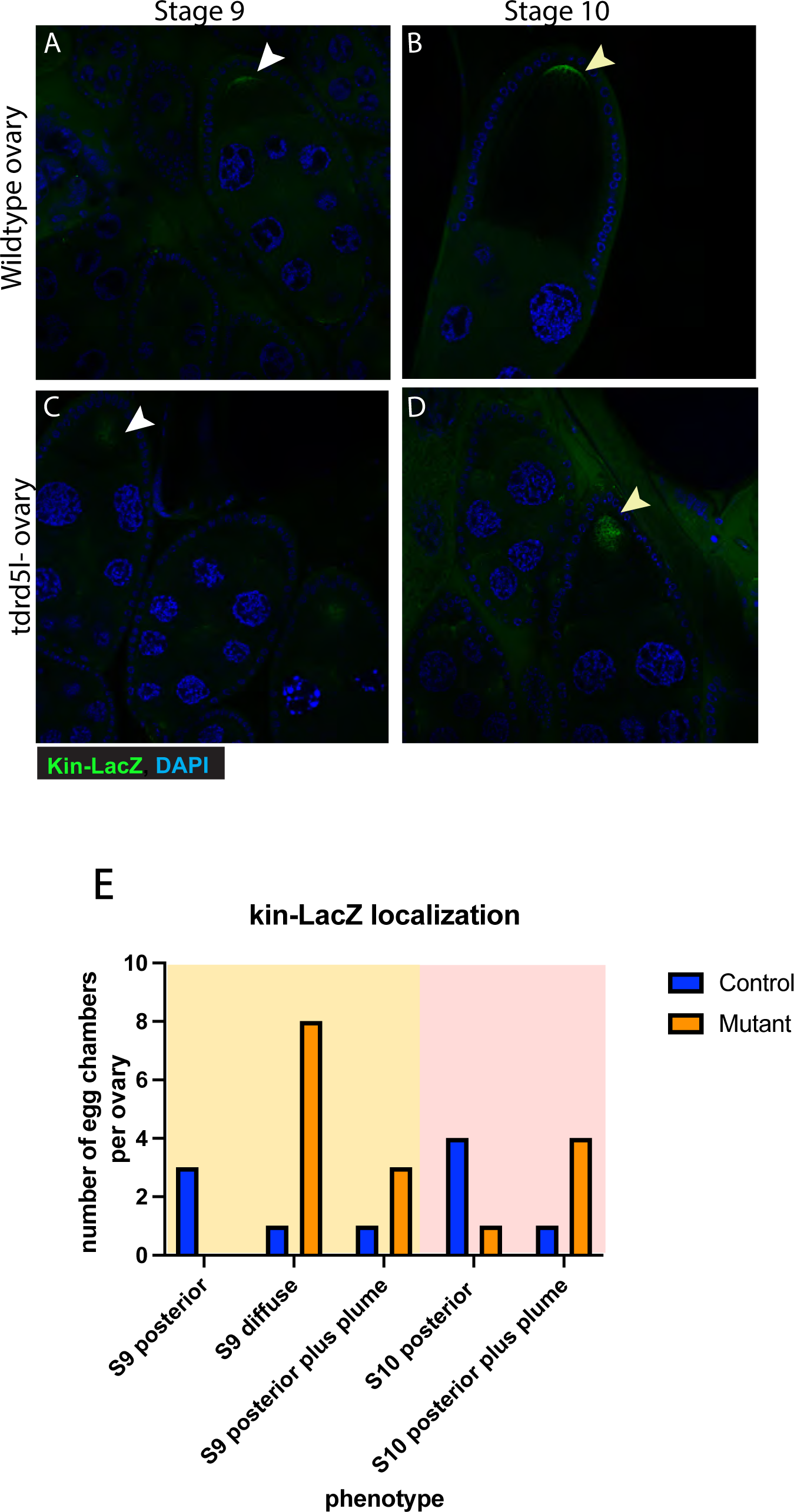
Kinesin mis-localizes in *Tdrd5l*-mutant oocytes. (A-D) Immunofluorescence of a Kin-LacZ reporter in developing ooctyes. Kin-LacZ expression is localized to the poster of the oocyte in wildtype (Kin-LacZ) stage 9 egg chambers (A) as marked by a white arrowhead and in wildtype (Kin-LacZ) stage 10 egg chambers (B) as marked by a yellow arrowhead. Kin-LacZ expression is not tightly localized to the posterior of the oocyte in *Tdrd5l*-mutant (*Tdrd5l^ΔM4/ΔQ5^; Kin-LacZ*) stage 9 egg chambers (C) as marked by a white arrowhead, or in *Tdrd5l*-mutant (*Tdrd5l^ΔM4/ΔQ5^; Kin-LacZ*) stage 10 egg chambers (D) as marked by a yellow arrowhead. (E) Quantification of observations in A-D. Quantifications of stage 9 egg chambers are in the yellow portion of the graph. Quantifications of stage 10 egg chambers are in the pink portion of the graph. *Tdrd5l* mutants are represented by orange bars, and controls are represented by blue bars. Each bar represents the percentage of egg chambers analyzed for that stage. N is noted inside each bar.

**Supplemental table 1:** Male RNAseq.

**Supplemental table 2:** Female RNAseq.

